# MXP: Modular eXpandable framework for building bioinformatics Pipelines

**DOI:** 10.1101/329110

**Authors:** Mikhail Kovtun, Igor Akushevich, Konstantin Arbeev, Anatoliy Yashin

## Abstract

**Background:** Pipelines are a natural tool in bioinformatics applications. Virtually any meaningful processing of biological data involves the execution of multiple software tools, and this execution must be arranged in a coherent manner. Many tools for the building of pipelines were developed over time and used to facilitate work with increasing volume of bioinformatics data. Here we present a flexible and expandable framework for building pipelines, MXP, which we hope will find its own niche in bioinformatics applications.

**Results:** We developed MXP and tested it on various tasks in our organization, primarily for building pipelines for GWAS (Genome-Wide Association Studies) and post-GWAS analysis. It was proven to be sufficiently flexible and useful.

**Conclusions:** MXP implements a number of novel features which, from our point of view, make it to be more suitable and more convenient for building bioinformatics pipelines.

## Background

MXP is a tool developed with intention to allow one to build pipelines easily.

MXP core (called “MXP base” below) is a set of Bash scripts that arrange execution of other scripts, called “methods”. This arranged execution is a pipeline. Drawing the analogy between MXP and languages like Python or R, MXP core corresponds to language interpreter, groups of methods correspond to packages, and pipelines correspond to end-user applications.

Pipelines may be very general or very specific, as any program can be. A distinguishing feature of MXP is that it allows you to easily modify or extend existing pipelines without changing the original pipeline code.

Two other distinguishing MXP features are using directories as units which pipelines operate on and the way to decide whether a target is up-to-date or should be rebuilt. These problems are significant in bioinformatics, and all tools for building pipelines have to struggle with them. MXP presents a novel approach to these problems.

In terminology of [1], MXP is an implicit configuration-based framework with a command-line interface.

## Implementation

### Approach

An important decision that should be made at the very beginning is what are units which framework operates on. In bioinformatics, the unit rarely is a single file. Much more often it is group of files, and sometimes very complex groups of files. For example, even alignment of paired-end reads requires to specify 2 input files; PLINK normally uses a triplet of files (`.bed, .bim, .fam`), and often it should be accompanied with files specifying set of SNPs to work with, phenotypic files, etc.

For these reasons, MXP uses filesystem directories as units. A directory can accommodate virtually any file structure. Additionally, it provides an easy solution for problems of where to store and how to find support files needed for the framework itself and helpful to the user (e.g., configuration that was used to obtain a target, log files, etc.).

Another important decision is to define the way to decide whether a target should be rebuilt. Often such a decision is made based on file timestamps: a target should be rebuilt if any required target is newer. This approach is inspired by make [2], and in the case of `make` it is very natural approach. However, in bioinformatics applications it is a rare case when input files are changed; instead, a user usually wants to change some parameters for an application (thresholds, window sizes, etc.) and re-run application with these new parameters. Detecting what targets are affected by such changes and therefore should be rebuilt is cumbersome.

To cope with this problem, MXP stores all parameters and scripts used to obtain a target, and checks whether they were changed in order to decide whether a target should be rebuilt. Our use of MXP demonstrated that the overhead caused by this approach is negligible.

MXP base is the core engine that executes a method’s scripts in the order prescribed by the Makefile.

MXP is written in pure Bash, and all units which are handled by MXP — methods, parameter sets, even Makefile — are Bash scripts. Of course, method scripts may invoke applications written in other languages, but any MXP-related script is still a Bash script.

This approach gives all power of the Bash to the pipeline writer. On the other hand, it has its own drawbacks, as Bash syntax is very cryptic and restrictive. However, we believe that the advantage of having the full power of Bash at hand outweighs the inconveniences.

### MXP overview and concepts

A pipeline is a sequence of operations that leads to a required result.

What exactly “result” means, and what kind of “operations” are used, depends heavily on the application domain. The expected application domain influences the design of a tool for building pipelines.

The famous Unix utility make [2], known since 1976, was, probably, the first tool for building pipelines (although the word “pipeline” is rarely used in conjunction with make). Virtually all of the tools for building pipelines borrow from make, and MXP is not an exception. But what makes these tools different are the elementary units which the pipeline operates on, how steps of pipeline are described, and the rules that are used to determine whether to re-execute a step or to use its existing results. This difference eventually influences the language used to describe the pipelines (e.g., Makefile syntax and semantics).

The units which MXP operates on are called (just like in make) targets. A target is represented by a directory containing an arbitrary set of files (and possibly subdirectories). We often use the word “target” instead more exact term “target directory”.

As in case of make, the execution of MXP consists of obtaining target specified in the command line. In order to obtain a target, other target(s) may be needed. MXP checks whether the required targets have been already obtained and if they are up-to-date; if not, MXP automatically rebuilds the required targets — which may require other targets, i.e., this is a recursive process. What targets are required for a given target, and how a given target should be obtained from the other ones is specified in Makefile (again, the term is borrowed from make).

### What is Makefile and how to use it

Makefile consists of rules. In MXP, Makefile is a Bash script. Here is a simple example of a rule:

~~~
MXP_MAKEFILE[d01_pdata]=” \
    (idata_DIR = d00_idata) pdata_0: pdata”
~~~

It is a Bash statement. It assigns string `“ (idata_DIR = d00_idata) pdata_0: pdata”` to an entry in associative array `MXP_MAKEFILE` indexed by string `“d01_pdata"`.

This rule states that:

- target `d01_pdata` requires target `d00_idata`
- method pdata with parameters `pdata_0` should be used to obtain target `d01_pdata` from target `d00_idata`
- during execution of method `pdata` environmental variable idata_DIR will be set to a full path to target directory `d00_idata`

Also, it implicitly states that:

- there is an analysis directory (current directory or directory explicitly specified in MXP commandline arguments) that contains a subdirectory mxp, and a file `Makefile.sh` inside of it
- the target directory named `d01_pdata` will be created within the analysis directory as a result of obtaining target `d01_pdata` (or, if this directory already exists, MXP will check whether this directory is up-to-date and rebuild it if it is not)
- there is a file pdata.sh containing a Bash script that will be executed in order to obtain target `d01_pdata`
- there is a file `pdata_0.params.sh` containing a Bash script (that define parameters) that will be executed in order to obtain the target `d01_pdata` Strictly speaking, there is no difference between a parameter script and a method script. MXP introduces this distinction to encourage the pipeline developers to clearly separate parameters from methods. Parameters could be changed by the pipeline user (for example, the user may want to use his own parameters for quality control), while methods are much more stable and are not expected to change from one pipeline application to another.

To determine if the target is up-to-date, MXP will check if:

- the target directory exists
- the last attempt to build target was completed successfully
- all required targets are up-to-date
- the rule used to obtain target has not been updated
- method and parameter scripts used to obtain target have not been updated

### Chaining pipelines

An important feature of MXP is that it allows for creating new pipelines by re-using pieces from existing pipelines. Each pipeline has a parent; only the root pipeline (which is a part of MXP base) does not have a parent. Makefile, methods and parameter sets defined in the parent pipeline are available in the child pipeline, and the child pipeline may override exactly those pieces from the parent pipeline that need to be changed.

In particular, the parent pipeline may be read-only, and still any fine-grained modifications of the parent pipeline are available to the user.

We anticipate that this feature will be widely used.

### Logging

Another important feature of MXP is logging. When a target is built, a full log is automatically written in the target directory. This log can be examined later to learn how exactly the target was built (in the case of successful build) or find out why the target build failed (in the case of failure).

It is also possible to save a log of a full MXP run, which may involve building multiple targets.

### Sharing and publishing pipelines

Reproducibility is very important for biological analyses, and, unfortunately, it is a weak point of many publications. We hope that MXP will significantly improve the ability of the researcher to publish information that describes exactly how results were obtained. To accomplish this, one needs to compress the mxp subdirectory of the analysis directory and submit the compressed file as a part of supplementary data.

## Results

We used MXP in our organization to build various pipelines. Primarily, we were interested in GWAS and post-GWAS analysis. MXP proved to be a convenient and easy-to-use tool for this purpose. We plan to publish an MXP-based GWAS pipeline as soon as it is finalized and documented.

Another application was the creation of a pipeline for obtaining and preprocessing files from public databases that are needed for annotating results of our analyses.

Using Bash as a programming language may seem to make the framework very slow. However, it is not the case. We use Bash carefully, and optimize all areas that may cause a slowdown. Running MXP when a target is already built (in this case MXP analyses hierarchy of Makefiles, checks that everything is up-to-date, reports it and terminates) takes about 1 second.

## Discussion

At the moment, multiple tools for building bioinformatics pipelines exist. [3] lists about 100 such tools. So, the question “why one more tool?” should be answered.

### The novel features

MXP has a few novel features (that up to our knowledge were not implemented in other tools). We already mentioned them in different contexts; here is the summary.

#### Directories as targets

Targets in MXP are represented by directories. It serves several purposes. First, it simplifies specification of methods’ input and output: when a method uses multiple input files and produces multiple output files, there is no need to specify all individual files explicitly (which is tricky when a set of input/output files is variable) – it is sufficient to specify output directory (i.e., the target being built) and input directories (i.e., the list of required targets). Second, it gives a simple answer to the question where to store supplementary files (i.e., files used by framework itself, logs, etc.). Third, it gives the user flexibility to combine several operations in one method (e.g., add reformatting the output of the main application of the method). This allows the user to reduce the number of targets, and make overall pipeline more manageable.

#### Comparing scripts to decide whether a target is up-to-date

In bioinformatics, tuning application parameters get correct result (e.g., quality control parameters may depend on dataset). When parameters for a target are changed, this target should be rebuilt, as well as all other targets that depend on it. To achieve this automatically, MXP stores all scripts used to obtain target in `.mxp` subdirectory of target directory. Then, when MXP checks whether a target is up-to-date, it compares the stored scripts with the current version of these scripts. If any difference is found, target will be rebuilt. The same effect may be achieved with other pipeline building tools, but it requires special work, while MXP does this automatically.

#### Ability to replace arbitrary piece of code without updating everything

Recall that every pipeline has a parent pipeline (except the root pipeline). When MXP needs to execute a script, it first searches the mxp subdirectory of the current analysis directory for this script. If not found, it searches parent pipeline, etc. Thus, if the user needs the parent pipeline with modification to a single script, he puts the modified script in mxp subdirectory of his analysis directory — and that is all what is needed.

The parent pipeline remains untouched (it may be read-only for the user). Other users who use the same parent pipeline are unaffected.

### The choice of languages

At least two languages are involved into construction of a tool for building pipelines: first, implementation language (which may be a combination of languages) and domain specific language (DSL), which is used to specify a pipeline. The better cooperation between these languages, the more convenient tool will be.

Python is often used as implementation language (Ruffus [4], Rubra [5], Omicspipe [6], Moa [7], pypeFLOW [8], PyPPL [9], Snakemake [10], and many other). Java and Groovy is another popular choice (BigDataScript [11], Bpipe [12], Nextflow [13], etc.). Occasionally, other languages like Prolog (Biomake [14]) or R (flowr [15]) were used.

Of course, languages like Python or Java have better syntax than Bash does and provide much more flexible data structures.

But at the very end pipeline should execute shell commands. Consequently, DSL contains lines (some-time quoted) that a shell commands. These commands necessary contain variables, which leads to a question who have to perform variable substitution: DSL implementation or shell? If DSL is chosen, the substitution is usually limited (no one is willing to implement the full analogue of Bash); if shell should perform substitution, the ability to communicate variable values to shell is a limiting factor.

For these reasons we chose Bash as the language to implement MXP. The only place where DSL is used in MXP is a rule for obtaining a target; i.e., the string value assigned to an entry in MXP_MAKEFILE associative array is a DSL statement. Makefile as a whole is a Bash script. Using Bash gives MXP several advantages. First, MXP may provide (and it does) convenience Bash functions that can be used in scripts implementing methods. Second, Bash arrays may be passed from parameter scripts to method scripts (as parameter and method scripts are sourced—rather than executed—in Bash subshell). Third, as Makefile is a Bash script, it may use all Bash features to create rules: for example, many similar rules may be generated in a simple Bash loop.

### MXP versus other tools

First, let us note that virtually any tool can be successfully used to build virtually any pipeline. For example, [16] demonstrates that even make can be used for bioinformatics pipelines. The question is convenience for specific applications.

MXP shares many features with other frameworks. We describe its distinguishing features above; from our point of view, they may provide MXP its own niche.

## Conclusions

MXP is a tool for creating pipelines, and therefore may be useful for researchers who are knowledgeable in programming are willing to create their own pipelines.

Our goal is to create reusable pipelines, primarily in the domain of GWAS and post-GWAS analyses. This work is similar to the one done in Omicspipe [6], which extends Ruffus [4] to create pipelines for analysis of results of Next Generation Sequencing (NGS). For our purpose, we consider MXP as more suitable for out tasks tool.

MXP is a stable and ready to use software. On the other hand, it is under development, and new features are added.

## Availability and requirements

**Project name**: MXP: Modular eXpandable framework for building bioinformatics Pipelines

**Project home page:** https://sites.duke.edu/barusoftware/MXP

**Operating system:** Linux/Unix

**Programming language:** Bash

**Other requirements:** Bash v.4.0 or higher

**License:** MIT (https://opensource.org/licenses/MIT)

## List of abbreviations

DSL: Domain-Specific Language
GWAS: Genome-Wide Association Study
MXP: Modular eXpandable framework for building bioinformatics Pipelines
NGS: Next Generation Sequencing

## Ethics approval and consent to participate

Not applicable.

## Consent for publication

Not applicable.

## Availability of data and materials

Not applicable.

## Competing interests

The authors declare that they have no competing interests.

## Funding

Research reported in this publication was partly supported by the National Institute on Aging of the National Institutes of Health (NIA/NIH) under Award Numbers P01AG043352, R01AG046860, and P30AG034424. The content is solely the responsibility of the authors and does not necessarily represent the official views of the National Institutes of Health.

## Authors’ contributions

MK designed and implemented MXP core. IA and KA participated in discussions about MXP design and implemented a number of methods for GWAS and post-GWAS analysis. AY provided overall guidance.

## Acknowledgments

Not applicable.

